# The lipid droplet protein Pgc1 controls the subcellular distribution of phosphatidylglycerol

**DOI:** 10.1101/308874

**Authors:** Dominika Kubalová, Petra Veselá, Thuraya Awadová, Günther Daum, Jan Malínský, Mária Balážová

## Abstract

The biosynthesis of yeast phosphatidylglycerol (PG) takes place in the inner mitochondrial membrane. Outside mitochondria, the abundance of PG is low. Here we present evidence that the subcellular distribution of PG is maintained by locally controlled enzymatic activity of the PG-specific phospholipase, Pgc1. We document that the Pgc1 absence leads to spreading of PG over various cellular membranes. Fluorescently labeled Pgc1 protein strongly accumulates at the surface of lipid droplets (LD). We show, however, that LD are not only dispensable for Pgc1-mediated PG degradation, but even host no phospholipase activity of Pgc1. Our *in vitro* assays document the capability of LD-accumulated Pgc1 to degrade PG upon entry to membranes of the endoplasmic reticulum, mitochondria, and even of artificial phospholipid vesicles. FRAP analysis confirms continuous exchange of GFP-Pgc1 within individual LD *in situ*, suggesting that a steady-state equilibrium exists between LD and membranes to regulate immediate phospholipase activity of Pgc1. In this model, LD serve as storage place and shelter Pgc1 preventing untimely degradation, while both phospholipase activity and degradation of the enzyme occur in membranes.

## Introduction

Phosphatidylglycerol (PG) is an anionic phospholipid which exhibits a great variety of biological functions. It is one of the major structural lipids in bacterial membranes ((Sohlenkamp & Geiger, 2016), and citations therein). In eukaryotic cells, PG represents a minor fraction of total glycerophospholipds, but it plays indispensable roles in optimizing their energy metabolism. Moreover, PG is an essential metabolic precursor of cardiolipin, the main lipid of the inner mitochondrial membrane (Horvath & Daum, 2013; Zinser & Daum, 1995). Cells deficient in PG biosynthesis exhibit severe phenotypic defects related to impaired mitochondrial function (Chang *et al*, 1998a; Janitor *et al*, 1995; Zhong & Greenberg, 2005; Zhong *et al*, 2005, 2007). PG is also an important structural and functional lipid of photosynthetic membranes (Kobayashi, 2016). For example, *Arabidopsis* mutants deficient in PG synthesis fail in thylakoid membrane formation (Babiychuk *et al*, 2003; Hagio *et al*, 2002; Haselier *et al*, 2010; Kobayashi *et al*, 2015). Extracellular PG secreted by type II alveolar cells participates in regulation of innate immune response within the lung and in prevention of viral infections (Hallman *et al*, 1985; Kandasamy *et al*, 2011; Numata *et al*, 2010, 2012, 2013. And, a product of PG degradation, diacylglycerol, is an important signaling molecule (Roelants *et al*, 2017; Stahelin *et al*, 2004).

In yeast, PG is a low-abundant phospholipid. It is biosynthesized in the inner mitochondrial membrane (Schlame & Haldar, 1993) and either utilized as substrate by the cardiolipin synthase (Crd1) (Chang *et al*, 1998b; Jiang *et al*, 1997; Tuller *et al*, 1998) or degraded by a PG-specific phospholipase C (Pgc1) (Pokorná *et al*, 2015; Simocková *et al*, 2008). The cellular level of PG is subject of tight regulation, and its misbalance influences deleteriously various cell functions. Cells lacking the phosphatidylglycerolphosphate synthase Pgs1, an enzyme responsible for the rate-limiting step of PG synthesis, lack both PG and cardiolipin. These cells fail to grow on nonfermentable carbon sources, are heat-sensitive (Chang *et al*, 1998a), and their mitochondria contain extremely elongated cristae sheets, which frequently form onion-like structures (Connerth *et al*, 2012). Mutants accumulating PG exhibit weaker, but still significant phenotypes. Cells lacking Crd1 accumulate PG in the absence of cardiolipin. Accordingly, most of their abnormal features, such as compromised stability of respiratory supercomplexes and mitochondrial DNA or decreased mitochondrial potential, result rather from the absence of CL than from the excess of PG (Baile *et al*, 2014; Jiang *et al*, 2000; Luévano-Martínez *et al*, 2015; Pfeiffer *et al*, 2003; Zhang *et al*, 2002; Zhong *et al*, 2004). However, phenotype differences between *pgs1*Δ and *crd1*Δ strains, which both lack cardiolipin but contain different amounts of PG, suggest specific roles for PG as well. These roles could be directly identified in a *pgc1*Δ strain, which accumulates PG without significant changes in the other phospholipids (Simocková *et al*, 2008). Cells lacking the *PGC1* gene exhibit fragmented mitochondrial network and increased respiration rates due to elevated activity of cytochrome *c* oxidase (Pokorná *et al*, 2015).

The subcellular localization of the two enzymes depleting the limited cellular stocks of PG, Crdl and Pgcl, strongly differ. While Crd1 activity has been localized exclusively to the inner mitochondrial membrane (Schlame & Haldar, 1993), the distribution of Pgc1 protein was confined to lipid droplets (LD), as reported by cell-fractionation and fluorescence microscopy data (Beilharz *et al*, 2003; Currie *et al*, 2014; Grillitsch *et al*, 2011; Ruggiano *et al*, 2016; Simocková *et al*, 2008). LD are universally conserved dynamic organelles, which exhibit a unique topology consisting of a hydrophobic core, predominantly composed of triacylglycerol and sterol esters, coated by a phospholipid monolayer and a set of specific proteins. The primary functions of LD are the storage of lipids and protection against lipotoxicity (Garbarino *et al*, 2009; Valachovic *et al*, 2016; Korber *et al*, 2017). However, more functions of LD were discovered, including an involvement in cellular signaling, lipid trafficking, protein storage, autophagy, and sporulation (Barbosa *et al*, 2015; Daum *et al*, 2007; Hsu *et al*, 2016; Leber *et al*, 1998; Singh *et al*, 2009; Wang, 2014, 2015).

In this study, we address the role of LD in the homeostasis of PG – the anionic lipid synthesized in mitochondria. We show that the absence of the LD protein, PG specific phospholipase Pgc1, causes spreading of PG over various cellular membranes. Together with *in vitro* tests of Pgc1 activity, our data suggest that the enzyme is inactive at the surface of LD and gets activated only after the translocation to intracellular membranes. Consequently not LD, but rather membranes of the neighboring organelles (endoplasmic reticulum (ER) and mitochondria) host the PG degradation. In this model, LD serve as storage places for the Pgc1 enzyme allowing for effective regulation of PG levels.

## Results

### PG-specific phospholipase Pgcl localizes exclusively to LD

Various analyses localized Pgc1 to LD (Beilharz *et al*, 2003; Simocková *et al*, 2008; Grillitsch *et al*, 2011; Currie *et al*, 2014), and mitochondria (Sickmann *et al*, 2003). Fluorescently tagged Pgc1 was shown to accumulate exclusively in LD when yeast cells were grown in the rich medium (Ruggiano *et al*, 2016). We checked to what extent this localization pattern of Pgc1 depends on the levels of the Pgc1 substrate, PG. To visualize the subcellular distribution of the Pgc1 protein, we prepared strains expressing a GFP-tagged version of Pgc1. The specific phenotype of a *pgc1*Δ mutant, namely accumulation of increased cellular PG levels in inositol-free media (Simocková et al., 2008), was successfully complemented by the expression of GFP-Pgc1, indicating the functionality of the fusion protein (Fig EV1).

Inositol is a known repressor of PG biosynthesis (He & Greenberg, 2004; Zhong & Greenberg, 2003). Even in inositol-free medium, however, the vast majority of the GFP-Pgc1 fluorescence signal in *pgc1*Δ cells expressing the GFP-Pgc1 fusion was confined to the surface of LD (Fig 1). Non-uniformly sized bright spots or empty circles of the GFP-Pgc1 fluorescence, as visualized in individual confocal sections, were unequivocally identified with LD using the specific lipophilic marker LD540 (Fig 2A-D). FRAP analysis revealed a stable character of this distribution: only slow recovery of the fluorescence intensity with the half-time of 704±21 s was observed after photobleaching of the GFP-Pgc1 fluorescence signal corresponding to an individual LD (Fig 2E). The quasi-linear character of the recovery curve indicated a limited communication between individual LD.

**Figure 1.**
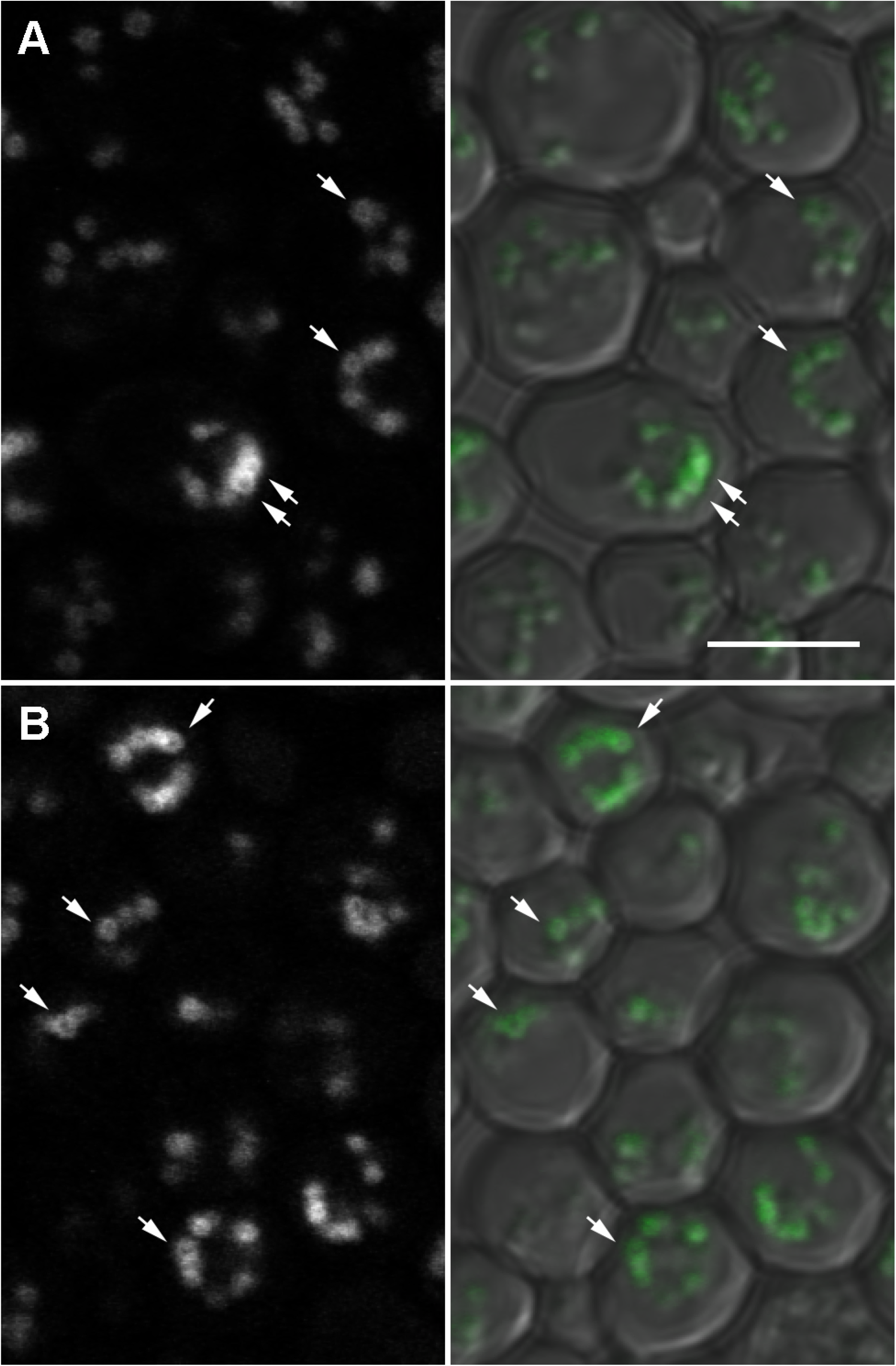
Localization of GFP-Pgcl protein is independent on induction of PG biosynthesis by exogenous inositol. A, B Cells expressing GFP-Pgc1 protein from pCM189 plasmid (strain *pgc1*Δ+*GFP-PGC1*, Table 1) were cultivated in SMD I+ (A) or I- (B) media without uracil. GFP fluorescence on single confocal sections (left) and composite images (GFP + DIC; right) are presented. Note that GFP-Pgc1 is localized on the surface of the labeled structures (arrows). Bar: 5μm.

**Figure 2.**
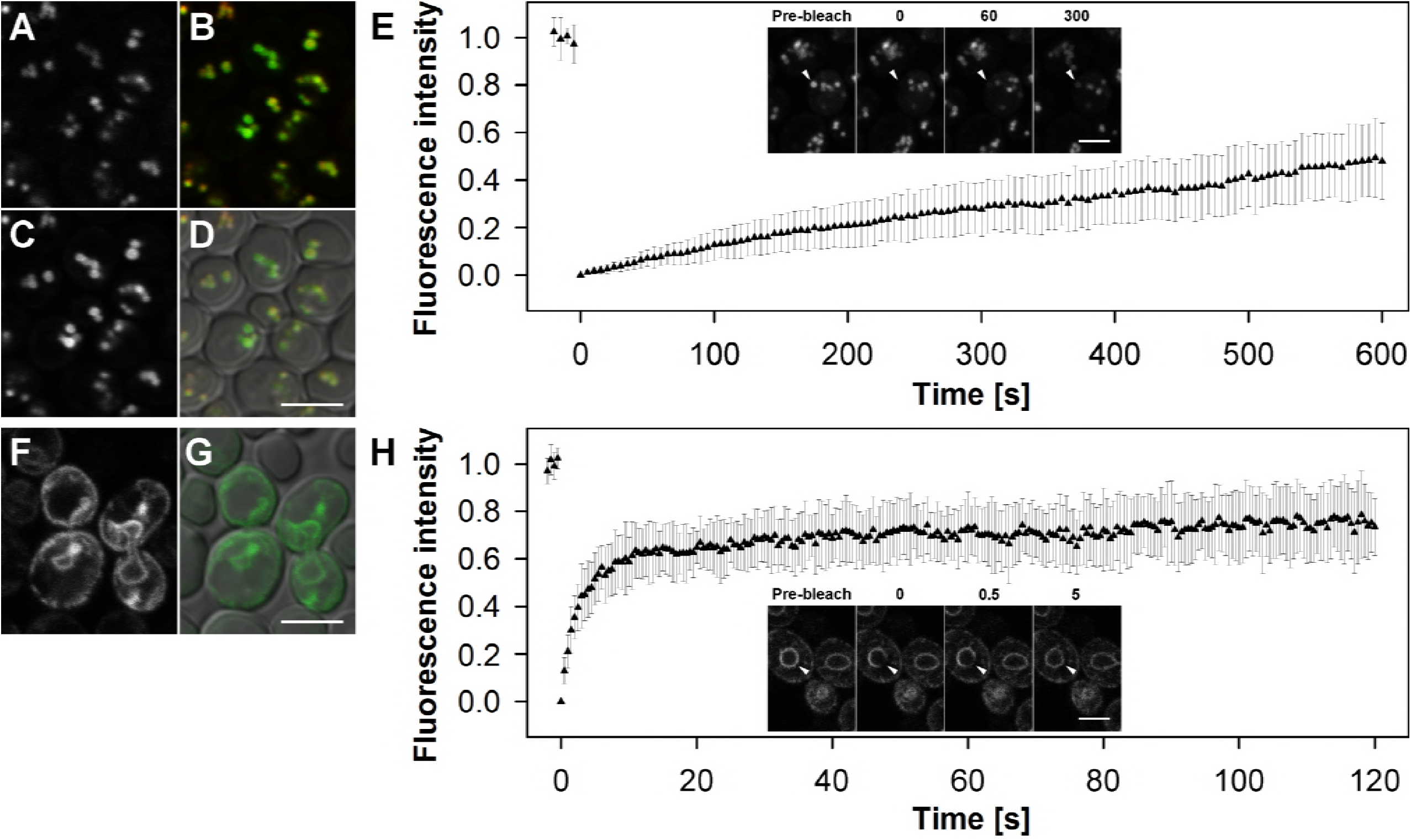
GFP-Pgc1 is stably sequestered to LD. A-E Cells expressing GFP-Pgc1 in the presence of LD (strain *pgc1*Δ+*GFP-PGC1*). F-H Cells expressing GFP-Pgc1 in the absence of LD (strain *QMpgc1*Δ+*GFP-PGC1*). E, H Mobility of GFP-Pgc1 was assessed using FRAP analysis. Relative fluorescence intensity changes monitored within the bleached area are plotted with error bars indicating SD (n=20). Inserts depict the distribution of GFP fluorescence before (Pre-bleach), instantly after the bleaching (0) and during the recovery (times indicated in seconds after the bleaching). Bar: 5 μm. Lipid droplet dye LD540 (A, red in B, D) and GFP fluorescence (C, E, F, H, green in B, D, G) on single confocal sections are presented.

A similar experiment was performed in the *pgc1*Δ mutant of a strain defective in LD formation (*dga1*Δ*lro1*Δ*are1*Δ*are2*Δ quadruple mutant strain, QM) (Sandager *et al*, 2002; Sorger *et al*, 2004; Athenstaedt, 2011). This scenario resulted in a completely different localization pattern of GFP-Pgc1. In QM*pgc1*Δ cells, the entire signal of the GFP-Pgc1 fluorescence was evenly distributed in membranes of the ER (Fig 2F, G). In contrast to GFP-Pgc1 localized in LD in the *pgc1*Δ strain, GFP-Pgc1 at perinuclear and cortical ER of the QM*pgc1*Δ cells was highly mobile. Fluorescence recovery half-time was estimated to 3.1±1.2 s, with the mobile fraction of 71±9% (Fig 2H).

### Deletion of the PGC1 gene causes accumulation of specific PG pools in various organelles

To address the subcellular localization of Pgc1 phospholipase activity *in situ*, we monitored the changes in the lipid composition of internal membranes caused by the deletion of *PGC1*. We expected PG to accumulate in those membranes of the *pgc1*Δ mutant, which in wild type strain represent either the places where the PG degradation is localized or at least places which communicate with places of PG degradation. Another strain showing the elevated levels of PG, *crd1*Δ, was used as a control to verify that the changes observed in *pgc1*Δ cells represented specific consequences of the *PGC1* deletion.

First, we analyzed the relative PG abundance in LD, mitochondrial and ER fractions of the wild type, *pgc1*Δ and *crd1*Δ cells (see Materials and Methods for details). Compared to the wild type, the relative abundance of PG was increased in all the analyzed subcellular fractions of the *pgc1*Δ mutant. No significant difference among LD, mitochondrial and ER membrane fraction could be seen; in all the fractions, PG levels represented ~4.5% of the total phospholipids (Fig 3A). In contrast, accumulation of PG in *crd1*Δ cells occurred almost exclusively in mitochondria, which exhibited a PG level of 8.6±1.1% of total phospholipids, while levels below 1% were detected in the LD and ER fractions of *crd1*Δ membranes.

**Figure 3.**
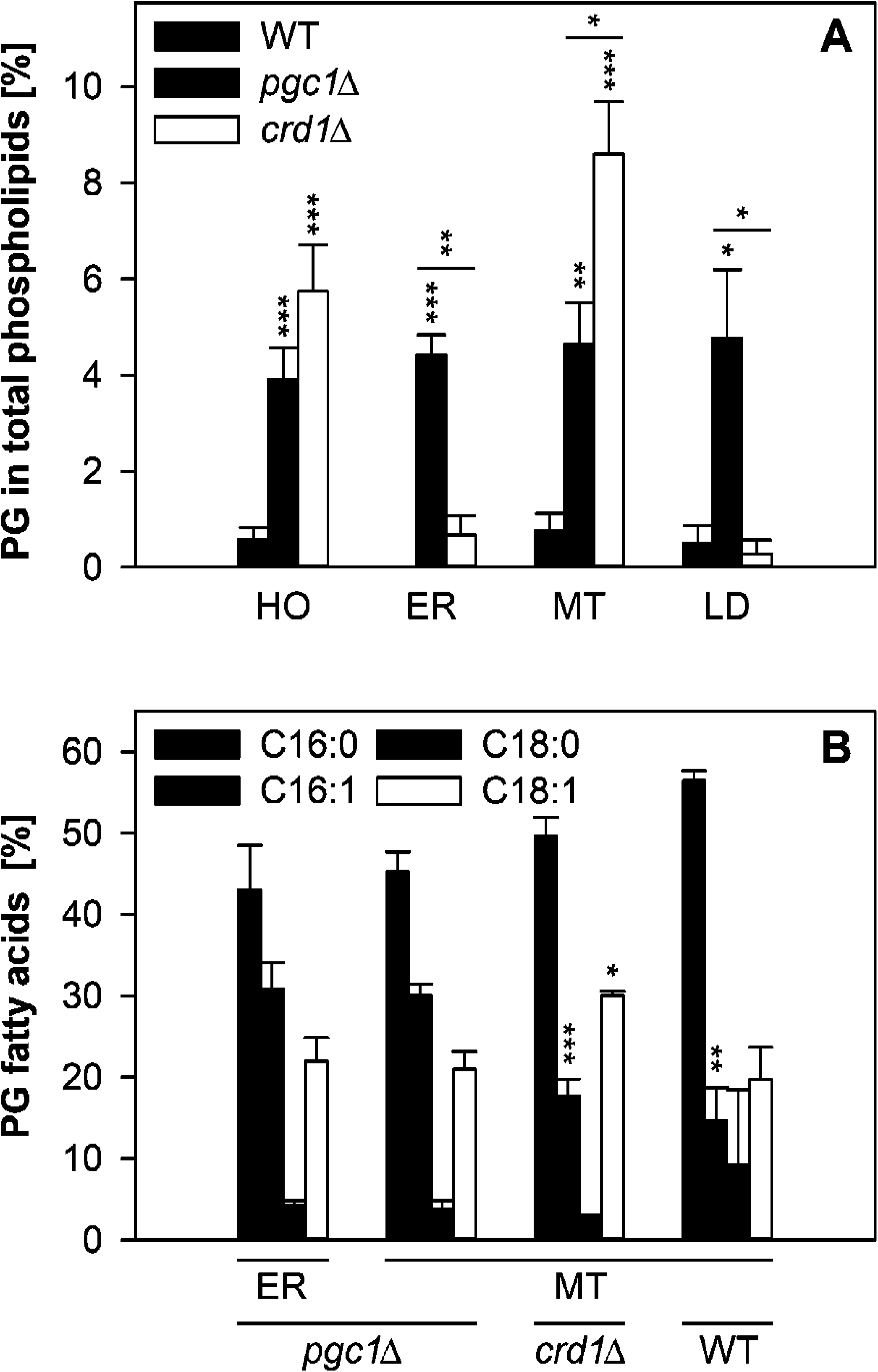
PG of specific fatty acid composition is evenly distributed in membranes of *pgcl*Δ cells. A In each fraction of wild type, *pgc1*Δ and *crd1*Δ cells, relative amount of PG (in % of total phospholipids) was evaluated. B PG fatty acids were estimated in mitochondrial fractions of all three analyzed strains, as well as in the ER fraction of *pgcl*Δ cells, by the gas chromatography. Relative amounts of individual fatty acids in respective fractions are shown. Mean values of at least four independent experiments ± SEM are presented. Statistically significant differences (see Materials and Methods for details) are marked by asterisks: * - p < 0.05; ** - p < 0.01; *** - p < 0.001. Data information: MT, mitochondria; ER, endoplasmic reticulum; LD, lipid droplets; WT, wild type.

Next, we compared the fatty acid composition of PG in the mitochondrial and the ER fraction of *pgc1*Δ. As shown in Fig 3B, no detectable difference between the two fractions could be identified, which indicated free PG exchange between ER and mitochondria. Both samples exhibited a significantly increased fraction of palmitoleic acid compared to the PG extracted from wild type mitochondria. Importantly, increased oleate instead of palmitoleic acid level was observed in mitochondrial membranes of *crd1*Δ cells. It is noteworthy that the acyl chain composition itself does not determine the Pgc1 phospholipase selectivity. As we showed previously, PG accumulated in mitochondrial membranes of *crd1*Δ cells could not be hydrolyzed by Pgc1 *in situ*, but it was accepted as a Pgc1 substrate *in vitro* (Pokorná *et al*, 2015). Altogether, these results could not indicate a specific localization of Pgc1 activity – all the analyzed fractions of *pgc1*Δ membranes exhibited similar abundance and acyl chain composition of the accumulated PG.

### Localization to a lipid bilayer is prerequisite for the enzymatic activity of Pgc1 protein

To directly test which organelle hosts the PG degradation, we analyzed *in vitro* activity of Pgc1 phospholipase in isolated membrane fractions. LD, mitochondrial and ER fractions were incubated with a fluorescent Pgc1 substrate (NBD-PG), and hydrolysis of this substrate to a fluorescent product (NBD-DAG) with altered thin layer chromatography mobility was used to measure Pgc1 activity in each fraction (Simocková *et al*, 2008).

In most of the analyzed strains, we observed ~50% hydrolysis of NBD-PG by Pgc1 in cell homogenates (Fig 4A). Comparable Pgc1 activities detected in wild type and QM mutant cells indicated that the presence of LD, the compartment accumulating the vast majority of Pgc1 in the wild type cells (Fig 2A-D), was not critical for PG degradation. This observation was further supported by the fact that the QM cells lacking LD did not exhibit elevated PG levels compared to the wild type (Fig EV2). Only low levels of NBD-PG hydrolysis were detected in all three analyzed sub-cellular fractions of wild type cells. Surprisingly, the highest NBD-DAG signal was detected in the ER fraction, while the Pgc1-enriched LD fraction of wild type cells exhibited no significant *in vitro* phospholipase activity.

Ruggiano and co-workers reported decreased degradation of Pgc1 protein in the strain lacking ubiquitin-protein ligase Doa10 (Ruggiano *et al*, 2016). Accordingly, we detected higher levels of Pgc1 activity in both ER and mitochondrial membrane fractions of *doa10*Δ strain compared to wild type. We observed similar increase of Pgc1 activity in a mutant strain lacking the mitochondrial outer membrane protein Msp1 that is also involved in protein quality control (Chen *et al*, 2014; Okreglak & Walter, 2014). However, the highest Pgc1 activity was detected in QM cells where all Pgc1 localized to membranes (predominantly to the ER membrane, Fig 2F, G). High Pgc1 activity was observed in both ER and mitochondrial fractions of QM cells, which indicated that not only the substrate (PG), but also the enzyme (Pgc1) can be exchanged between these two compartments. As apparent from the control experiments performed in *pgc1*Δ, *doa10*Δ*pgc1*Δ, *msp1*Δ*pgc1*Δ and QM*pgc1*Δ strains, the hydrolysis of NBD-PG to NBD-DAG could be ascribed predominantly to the activity of Pgc1 (Fig 4A).

**Figure 4.**
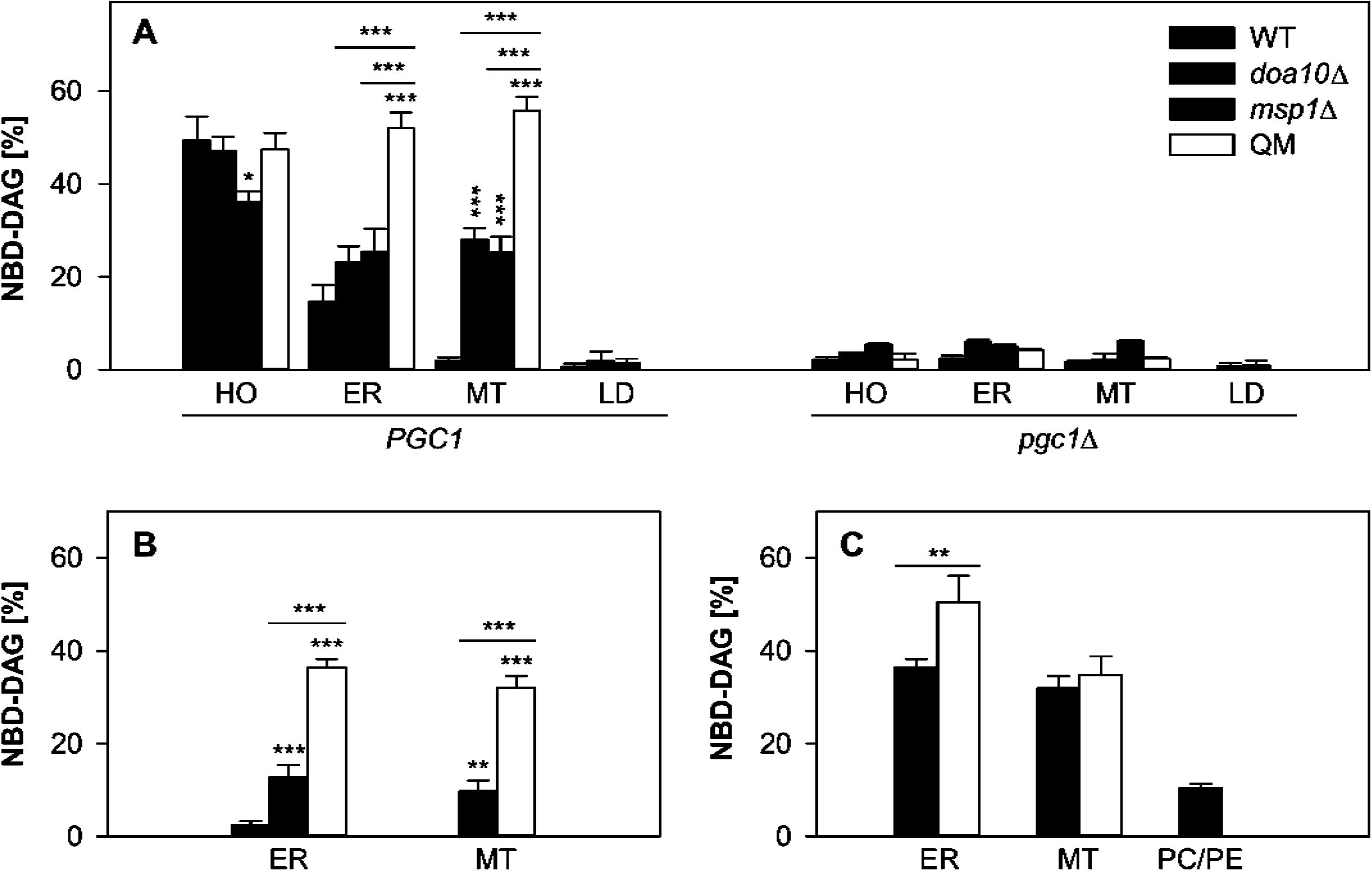
Pgc1 is active whenever embedded into a lipid bilayer. A Wild type, *doa10Δ, msp1*Δ, and QM cells with (*PGC1*) or without *PGC1* gene (*pgc1*Δ) were grown in SMD I- media for 24 h. *In vitro* phospholipase activity of Pgc1 in isolated subcellular fractions (HO, cell homogenate; ER, endoplasmic reticulum; MT, mitochondria; LD, lipid droplets) was measured in terms of relative amounts of NBD-DAG, a product of Pgc1-mediated hydrolysis of NBD-PG. Mean values of at least three and two independent experiments ± SEM are presented for *PGC1* and *pgc1*Δ strains, respectively. B LD with increasing amounts of Pgc1 protein were isolated from strains *pgc1* Δ, WT, and WT + *PGC1. In vitro* activity of Pgc1 in mixtures of these LD with the ER or mitochondrial (MT) fraction isolated from *pgc1*Δ strain was analyzed. C *In vitro* activities of Pgc1 in mixtures of LD isolated from the strain overexpressing *PGC1* (WT + *PGC1*) with membrane bilayers of different origin were analyzed. Mixtures containing complete ER and mitochondrial (MT) membrane fractions (black), membranes composed of isolated lipids from these fractions (white), and membranes composed of synthetic phospholipids (PC/PE; grey) are compared. Data information: In (B, C), mean values of at least four independent experiments ± SEM are presented. In A-C, Statistically significant (see Materials and Methods for details) differences are marked by asterisks: * - p < 0.05; ** - p < 0.01; *** - p < 0.001.

It is worth mentioning that the observed absence of Pgc1 activity in LD did not reflect the absence of the substrate – LD of *pgc1*Δ cells accumulated PG at levels similar to ER and mitochondrial membranes (Fig 3). Similarly, when NBD-PG was added to the LD fraction, it also massively associated with LD (Fig EV3). As an alternative explanation for the negligible NBD-PG hydrolysis in LD fraction of the wild type cells, we tested the hypothesis that Pgc1 activation required embedding of the enzyme into the phospholipid bilayer.

To confirm that the inability of Pgc1 to hydrolyze PG in the phospholipid monolayer of the LD surface (Figs 1, 2) can explain the results presented in Fig 4A, we measured *in vitro* Pgc1 activity in mixtures of the LD fraction with different membranes. Three types of Pgc1-less membranes were used: ER or mitochondrial fractions isolated from *pgc1*Δ cells, liposomes prepared from lipids isolated from these two fractions of *pgc1*Δ cells, and liposomes prepared from synthetic phospholipids (PC/PE, 3:1). In any of these membrane samples, no measurable spontaneous hydrolysis of NBD-PG could be detected. After the addition of the LD fraction isolated from wild type cells that contained Pgc1, we observed a clear increase of the NBD-PG hydrolysis product, NBD-DAG. This increase strongly depended on the Pgc1 protein content in the added LD fraction (Fig 4B). In LD-containing mixtures, either with complete ER or mitochondrial membrane fractions or with reconstituted bilayers from the respective extracted lipids, comparable amounts of NBD-DAG were detected. Lower, but significant NBD-PG hydrolysis was detected even in a mixture with liposomes prepared from synthetic lipids (Fig 4C). Apparently, the presence of a phospholipid bilayer was sufficient to induce the phospholipase C activity of Pgc1.

Another support for the hypothesis that the phospholipase activity of Pgc1 required the enzyme embedded into a lipid bilayer was provided by the analysis of *in vitro* Pgc1 activity in cell homogenates prepared from wild type and QM strains. For each strain, we compared Pgc1 activities in homogenates prepared by two different approaches. In the first approach, the cells were disintegrated with glass beads. This approach results in a high fragmentation of all cellular contents, including membranes. The second, gentler approach represented enzymatic cell wall digestion by zymolyase followed by breaking the cells in a Dounce homogenizer. In this case, the membrane fragmentation is significantly lower, this approach even preserves mitochondrial membrane respiration (Pokorná *et al*, 2015). Our data clearly indicated that the *in vitro* Pgc1 activity was significantly higher in gently prepared cell homogenates, which contained better preserved membranes. The detected difference in Pgc1 activity was about 2-fold in wild type, and about 5-fold in QM mutant preparations (Fig EV4). Concerning the difference between wild type and QM samples homogenized with beads, relatively higher Pgc1 activity detected in wild type cells could indicate an unspecific spreading of the Pgc1 protein from LD disintegrated during the treatment.

Since Ruggiano et al. (2016) suggested that the C-terminal domain of Pgc1 could be responsible for the membrane association of the enzyme, we tested localization and phospholipase activity of the C-terminally truncated Pgc1 proteins, Pgc1^1-274^ and Pgc1^1-297^. The expression level of Pgc1 constructs from plasmid was confirmed by RT-qPCR of the specific mRNA, and it was about 50fold higher compared to the expression of genomic *PGC1* in the wild type (Fig 5A). In the case of GFP-Pgc1 proteins, Western blot quantification of the protein level was performed. In *pgc1*Δand QM*pgc1*Δ cells, it showed low levels of both C-terminal truncated proteins GFP-Pgc1^1-274^ and GFP-Pgc1^1-297^ compared to full-length GFP-Pgc1 (Fig 5B). This corresponded well with our fluorescence microscopy observations: in cells expressing the GFP-Pgc1^1-274^ protein, just a faint GFP fluorescence homogenously dispersed in cytoplasm and none localized to LD could be detected (Fig 5C). As revealed by the *in vitro* NBD-PG assay, phospholipase C activities present in homogenates of cells expressing only truncated Pgc1 proteins, Pgc1^1-274^ and Pgc1^1-297^, were an order of magnitude lower compared to the full-length protein (Fig 5D). These reduced activities also reflect primarily low levels of truncated proteins in homogenates, although they also indicate the persisting enzymatic activity of the C-terminally truncated Pgc1 proteins. However, overexpression of none of the truncated proteins was able to fully complement the most prominent *pgc1*Δ phenotype – accumulation of PG (Fig 5E). Together with the aforementioned data, these results strongly supported our hypothesis that embedding of the Pgc1 protein into the phospholipid bilayer was prerequisite for its phospholipase activity.

**Figure 5.**
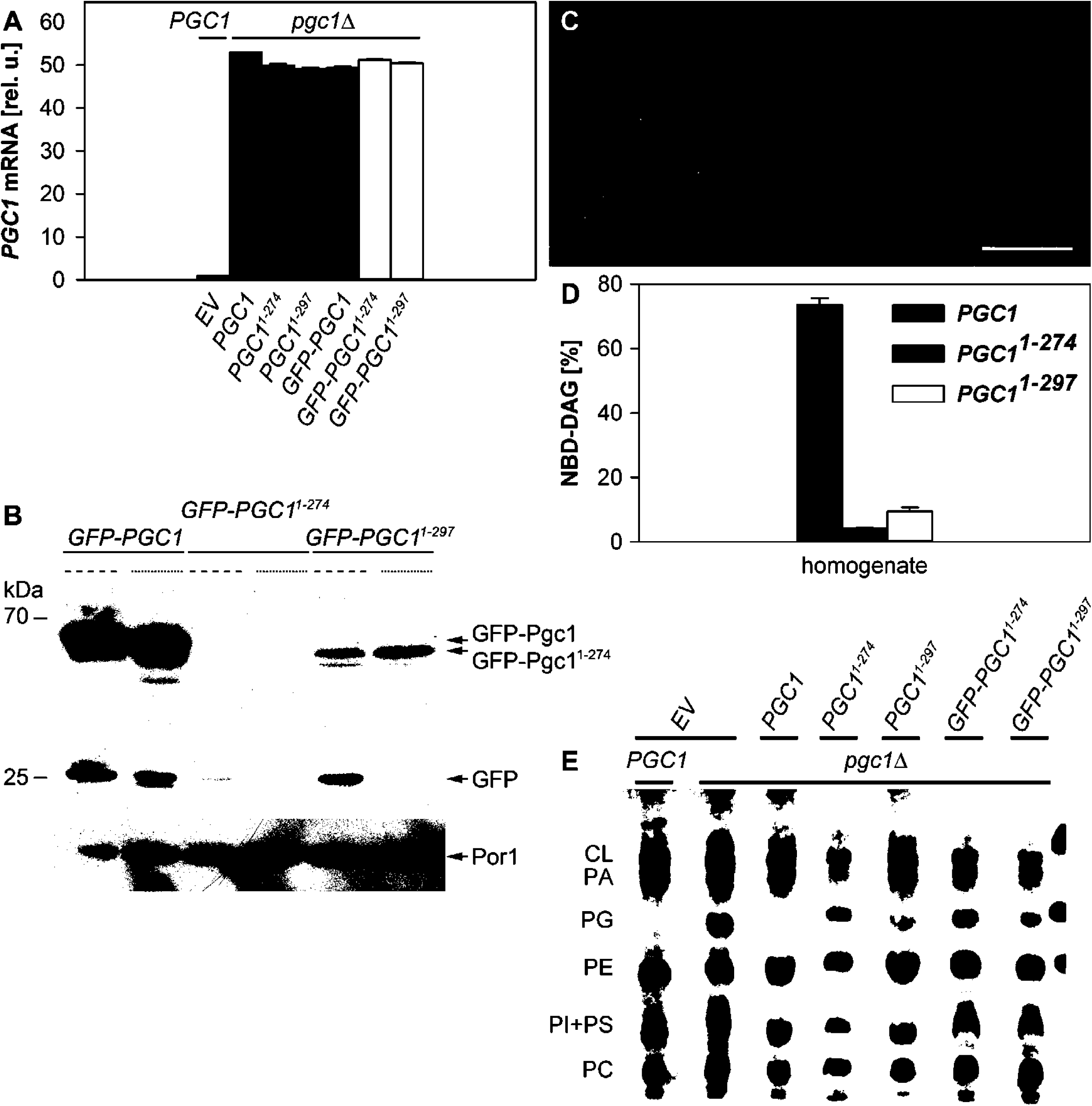
C-terminal truncation disrupts Pgcl function. A Levels of *PGC1, PGC1^1-274^*, and *PGC1^1-297^* mRNA in strains expressing the respective Pgcl protein versions either with or without GFP tagging on *pgc1*Δ background were compared. WT + EV strain was used as a control. The results were normalized to *ACT1* and *IPP1* mRNA levels; the wild type *PGC1* mRNA level was set to 1. Mean values of at least two independent experiments ± SEM are presented. B Using anti-GFP antibody, GFP-tagged Pgc1 versions were tracked on immunoblot. Por1 was used as a loading control (bottom panel). Protein levels in strains with (*pgclA*-derived; dashed line) and without LD (QM*pgc1*Δ-derived; dotted line) were compared. C GFP-Pgcl^1-274^ (strain *pgc1*Δ+*GFP-PGC1^1-274^*) was localized by fluorescence microscopy. A single confocal section is presented. Bar: 5 μm. D *In vitro* activities of Pgc1, Pgcl^1-274^ and Pgc1^1-297^ in cell homogenates (strains *pgc1*Δ+*PGC1, pgc1*Δ+*PGC1^1-274^* and *pgc1*Δ+*PGC1^1-297^*) were measured in terms of relative amounts of NBD-DAG, a product of Pgc1-mediated hydrolysis of NBD-PG. Mean values of at least three independent experiments ± SEM are presented. E Using thin layer chromatography, phospholipid profiles of strains overexpressing different Pgc1 versions were measured. Lipids extracted from cell homogenates were separated and visualized as described in Materials and Methods. PC, phosphatidylcholine; PI, phosphatidylinositol; PS, phosphatidylserine; PE, phosphatidylethanolamine; PG, phosphatidylglycerol; PA, phosphatidic acid; CL, cardiolipin; EV, empty vector.

## Discussion

While the biosynthesis of mitochondrial anionic phospholipids was clearly localized to the inner mitochondrial membrane (Schlame & Haldar, 1993; Dzugasová *et al*, 1998), little is known about subcellular localization of their degradation. Specifically, limited knowledge is available concerning the mechanisms by which relatively low concentrations of these lipids are maintained outside mitochondria. In this study, we tried to address this problem by the analysis of the enzymatic activity of a yeast PG-specific phospholipase C Pgc1 within the cell.

Under normal conditions, Pgc1 enzyme localizes exclusively to LD (Fig 1, see also (Simocková *et al*, 2008; Beilharz *et al*, 2003; Grillitsch *et al*, 2011; Ruggiano *et al*, 2016; Currie *et al*, 2014)), and as indicated by our FRAP data, its availability outside this compartment is strongly limited (Fig 2E). This localization pattern of Pgc1 is insensitive to the stimulation of PG biosynthesis in the absence of inositol (Fig 1). Even under the conditions of induced PG biosynthesis, LD-confined Pgc1 is capable to prevent PG accumulation in various locations – in the ER, mitochondrial and probably also in other membranes (Fig 3A). This fact indicates ongoing phospholipase activity of Pgc1 under these conditions.

The activity of Pgc1 however is independent on the LD compartment, as we detected comparable levels of PG degradation in wild type and QM strain, which is defective in LD formation (Fig EV2). Moreover, no phospholipase activity of Pgc1 could be detected in isolated Pgc1-containing LD fraction, suggesting that the Pgc1-mediated PG degradation occurs outside LD. Similar dual localization of the active and inactive enzyme has been documented before. Enzymatic activity of another LD protein, squalene epoxidase Erg1, takes place at the ER membrane. In contrast to Pgc1 however, partitioning of Erg1 to both LD and the ER membrane could be clearly visualized by fluorescence microscopy in wild type cells (Leber *et al*, 1998).

Not all PG outside LD is accessible for Pgc1-mediated degradation. For example, Pgc1 failed to remove PG accumulated in *crd1*Δ mutant. We suggested recently that this was probably due to the specific localization of the PG pool in *crd1*Δ cells (Pokorná *et al*, 2015). Data presented in Fig 3A strongly support this hypothesis, documenting a specific distribution of PG in *crd1*Δ cells. While rather homogenous spreading of PG over all analyzed cellular fractions could be observed in *pgc1*Δ cells, the *crd1*Δ mutant stored a high amount of PG only in mitochondria.

Moreover, PG accumulating in *crd1*Δ mutant exhibited a significant difference in acyl chain composition, if compared to PG accumulated by *pgc1*Δ cells (Fig 3B). This finding suggested that mitochondrial PG in *crd1*Δ cells could be a subject of remodeling, similar to CL in the wild type. Finally, PG accumulated by the *crd1*Δ mutant could be hydrolyzed by Pgc1 *in vitro* (Pokorná *et al*, 2015), indicating that not the acyl chain composition, but the localization of the lipid in these cells interferes with the Pgc1 function. We speculate that the observed difference in acyl chain composition, as well as enormous stability towards Pgc1-mediated degradation *in situ* (Pokorná *et al*, 2015) could reflect the fact that in *crd1*Δ cells, PG functionally substitutes the missing CL at the inner mitochondrial membrane.

Several lines of evidence indicate that the phospholipase activity of Pgc1 is not strictly determined by the composition of the surrounding membrane, but rather by the embedding of the protein into a lipid bilayer: i) while Pgc1 was inactive in pure LD fractions, it was capable to hydrolyze NBD-PG in our *in vitro* assay when a bilayer membrane was added to the reaction. This effect was similar whether biological membranes or just lipid membranes composed of isolated or synthetic lipids were used (Fig 4C); ii) phospholipase activity of Pgc1 in cell homogenates depended on the presence of intact membranes rather than on the original distribution of the Pgc1 protein *in situ* (Fig EV4); iii) phospholipase activity of C-terminally truncated Pgc1 proteins, which contained the whole glycerophosphodiester phosphodiesterase motif (Simocková *et al*, 2008), but lacked membrane localization domains, was effectively suppressed (Fig 5). Together these data suggest that the spatial sequestration of Pgc1 in a surface lipid monolayer of LD provides two independent ways to regulate the Pgc1 activity: 1) direct inhibition of the phospholipase function by keeping the protein outside the lipid bilayer, probably in a conformation which precludes its phospholipase activity, and 2) indirect inhibition by a strong limitation of the protein exchange between LD surface and membranes (Fig 2E).

Based on these data, we suggest a model of PG homeostasis mediated by the Pgc1 activity, which is tightly regulated by the localization and a rapid turnover of the enzyme (Fig 6). In this model, PG is generated at the inner mitochondrial membrane. The portion of the lipid which is not used for the CL biosynthesis freely spreads to other membranes within the cell along the concentration gradient. In these membranes, PG is cleaved by a specific phospholipase, Pgc1. Pgc1 seems to be a very effective enzyme, as just a minute in fluorescence microscopy invisible fraction of this protein is sufficient to keep PG at low level. But not lower than necessary: Pgc1 itself is a subject to fast degradation, executed by Ssm4/Doa10 ubiquitin-protein ligase at the ER membrane, and possibly by Msp1 AAA-ATPase at the outer mitochondrial membrane (Fig 4A; Ruggiano *et al*, 2016). Indeed, there are indications that Pgc1 can migrate to the outer mitochondrial membrane, at least under the conditions of overexpression (Simocková *et al*, 2008), or in the absence of LD (Fig 4A). Under normal conditions, LD provide a storage place for the vast majority of Pgc1. Localization to LD protects the protein against ubiquitination and the rapid turnover. Delivery of Pgc1 to the ER membrane is mediated by continuous LD-ER contacts; direct contacts of LD with other membranes are not *a priori* excluded either.

**Figure 6.**
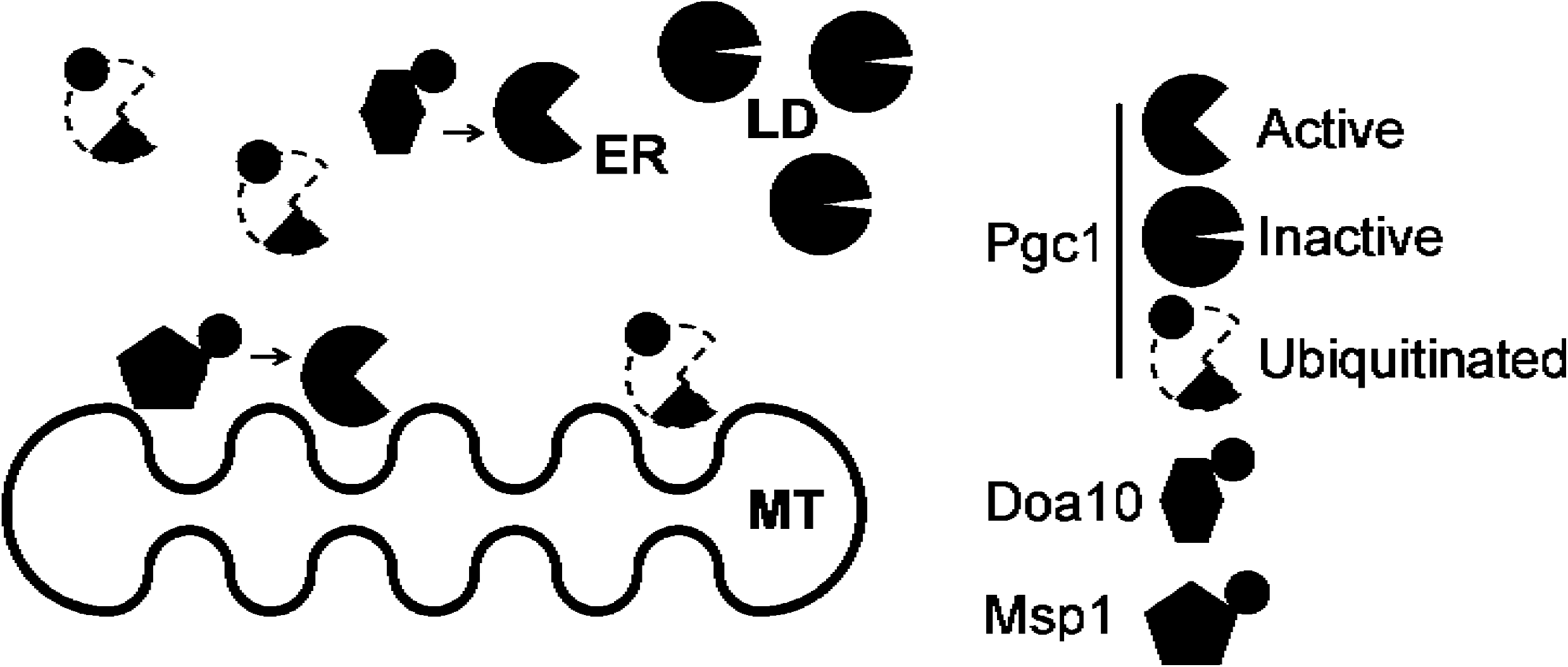
Model of PG homeostasis in *S. cerevisiae*. Yeast PG is synthesized at the inner mitochondrial (MT) membrane (dark grey) and freely spreads from there all over the cellular membrane system. In other membranes, its levels are kept low by the action of Pgc1. Inactive form of Pgc1 is stored at the surface of LD. Through the continuous contacts of LD with the ER membrane, Pgc1 diffuses to the ER and gets activated. Further spreading of the enzyme to other membranes is highly likely. Our data indicated Pgc1 activity in the outer mitochondrial membrane, for example. At membranes, Pgc1 becomes accessible for rapid ubiquitylation by local factors, such as Doa10 in the ER, and possibly Msp1 at the outer mitochondrial membrane. Actual PG levels are then a result of balanced i) PG synthesis, ii) Pgc1-mediated PG degradation triggered by the release of the ezyme from LD, and iii) Pgc1 turnover.

The described arrangement represents an effective mechanism capable of fast, wide-range, and bidirectional regulation of PG levels within the cell. What remains to be answered is the reason why the cells so tightly control the content of one minor phospholipid in their membranes, as apparently too low, as well as too high amounts of PG matter. So far, only a few indications are available in this respect. We know, for example, that the accumulation of PG compromises the barrier function of the inner mitochondrial membrane in *pgc1*Δ cells (Pokorná *et al*, 2015). In *C. elegans*, changes in mitochondrial morphology triggered after inactivation of CL synthase were recently also interpreted in terms of PG accumulation rather than lack of CL (Matsumura *et al*, 2018). Also, it is worth mentioning that the product of PG hydrolysis, DAG, is an important signaling molecule (Roelants *et al*, 2017; Stahelin *et al*, 2004). Further studies will be necessary to explain the specific roles of PG in the membrane system of eukaryotic cells.

## Materials and Methods

### Yeast strains and growth conditions

All yeast strains used in this study are listed in Table 1. Cultures were grown on complex media YPD (2 % yeast extract, 1 % peptone, 2 % glucose). For experiments, yeast were grown aerobically at 30 °C in a defined synthetic SMD medium prepared as described previously (Griac *et al*, 1996), with 2 % glucose as a carbon source. Synthetic medium was either supplemented with 75 μM inositol (I+) or lacked inositol (I-). Transformants were selected on synthetic medium without uracil or histidine.

**Table 1.**
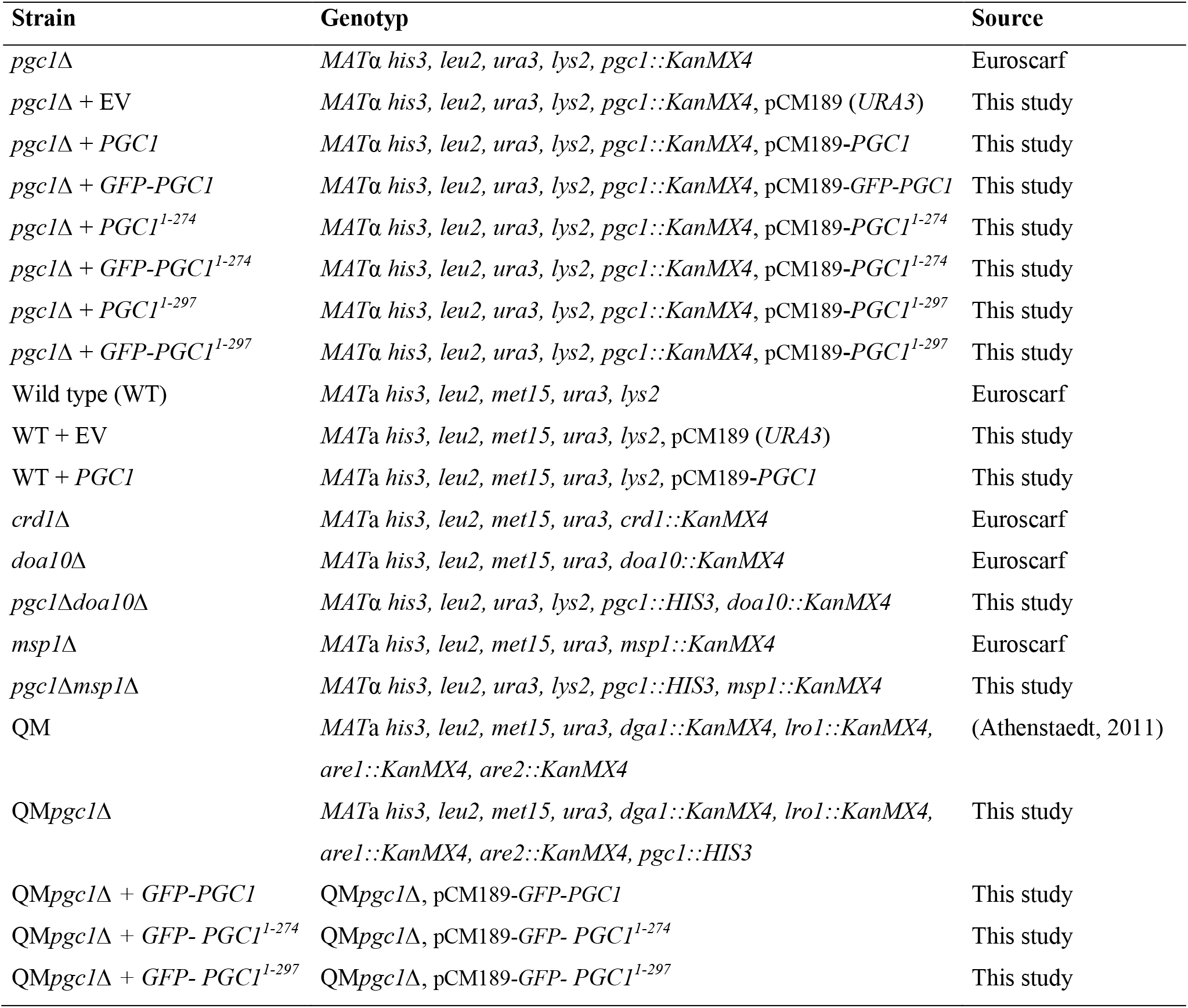
Yeast strains. All *S. cerevisiae* strains were in BY4741 or BY4742 background (Euroscarf).

### Plasmids and strains construction

Tetracycline-regulated centromeric expression vector pCM189 (Euroscarf, Scientific Research and Development GmbH, Germany) was used for the expression of *PGC1* genes. To prepare pCM189-*PGC1* plasmid, the *PGC1* gene was amplified by PCR from *S. cerevisiae* genomic DNA using 5’-ATCAGGATCCATGGTTGAAATTGTGGGCCAC-3’ and 5’- AGAGCTGCAGCAAAATATCACTATTCAAATCTCGC-3’primers. The PCR fragment was then ligated into pCM189 plasmid using *BamHI* and *PstI* restriction sites.

To prepare pCM189-*GFP-PGC1* plasmid, the *GFP* gene was amplified by PCR from the plasmid used before (Beilharz *et al*, 2003) using 5’-ATTGGATCCCGGGATGAGTAAAGGAGAAGAACTTTTCAC-3’ and 5’-ATTGGATCCCTGCAGTTTGTATAGTTCATCCATGCCAT-3’ primers. The amplified PCR fragment was digested with *BamHI* and cloned into the *BamHI* restriction site of the pCM189-*PGC1* plasmid. Both truncated versions of *PGC1* or *GFP-PGC1* were prepared by amplifying with the same forward 5’ primer, for *PGC1* versions (5’-ATCAGGATCCATGGTTGAAATTGTGGGCCAC-3’) and for *GFP-PGC1* versions 5’-ATTGGATCCCGGGATGAGTAAAGGAGAAGAACTTTTCAC-3’). Reverse primer 5’- ATTGCGGCCGCTCATACAGATGCCAGACTGGGCG-3’ was used for amplifying *PGC1^1-274^* or *GFP- PGC1^1-274^*, and 5’- ATTGCGGCCGCTCACCATTTGGAATATAGAAGCGT-3’ was used for amplifying *PGC1^1-297^* or *GFP- PGC1^1-297^* by PCR from pCM189-*GFP-PGC1*. The PCR fragments were then ligated into pCM189 plasmid using *BamHI* and *NotI* restriction sites.

Double mutants *pgc1*Δ*doa10*Δ and *pgc1*Δ*msp1*Δ were prepared as follows: a disruption cassette *pgc1*Δ::*HIS3* from *pgc1*Δ*crd1*Δ strain was PCR amplified (Simocková *et al*, 2008) and transformed into *doa10*Δ or *msp1*Δ strains, respectively. Similarly, the QM*pgc1*Δ strain was prepared. Yeast transformation was performed by the lithium acetate method (Gietz & Woods, 2002).

### Fluorescence Microscopy

Living yeast cells grown in SMD I- or SMD I+ medium were concentrated by brief centrifugation, immobilized on a 0.17 mm thick coverslip by a thin film of 1 % agarose in 50 mM phosphate buffer, pH 6.3, and observed using a confocal microscope LSM510 (Zeiss) with a 100× PlanApochromat oil-immersion objective (NA = 1.4). Fluorescence signals of GFP and LD540 (a generous gift of Dr. Christoph Thiele, Max Planck Institute of Molecular Cell Biology and Genetics, Dresden, Germany) were excited by 488 nm line of Ar laser and 561 nm line of solid state laser, and detected using bandpass 505–550 and 575–615 nm emission filters, respectively. For staining with LD540, the dye dissolved in ethanol was added to the cell culture (to final dye concentration of 0.5 μg/ml) and incubated in dark at the room temperature for 30 min.

### Fluorescence recovery after photobleaching

Fluorescence recovery after photobleaching (FRAP) experiments were performed in the single photon excitation mode on the laser scanning microscope FV1200MPE (Olympus) with a 100× PlanApochromat oil-immersion objective (NA = 1.4). Fluorescence signal of GFP was excited by 488 nm line of Ar laser, and detected using a bandpass 500–545 nm emission filter. A SIM scanner was used to minimize the time delay between the fluorescence photobleaching in the region of interest (ROI) and the acquisition of the first post-bleach frame. To quantify the initial level of fluorescence, five x-y frames were acquired prior to the bleaching. Circular ROI encompassing a whole individual LD or a part of the ER membrane, with a diameter of 1 μm, was bleached with 50 ms pulse of 405 nm laser at maximum intensity. The fluorescence recovery in LD and ER was monitored in 5 and 0.5 s intervals for 10 and 2 minutes, respectively. Acquired fluorescence intensities were corrected for the scan-induced photobleaching and normalized to the fluorescence intensity measured in a pre-bleach period. The curve fitting was performed in SigmaPlot 12.5 (Systat Software Inc.).

### Preparation of subcellular membranes

Mitochondrial and ER fractions were isolated by differential centrifugation according to Zinser and Daum (Zinser & Daum, 1995), with the following modifications: spheroplasts were prepared using Zymolyase 20T (Nacalai Tesque) from cells grown in SMD I- media for 24 h at 30 °C. The spheroplasts were homogenized in a tight-fitting Dounce homogenizer in 0.6 M mannitol; 10 mM Tris/HCl, pH 7.4; 1 mM phenylmethylsulfonyl fluoride. The homogenate was centrifuged for 5 min at 3400 g to remove residual cell debris. The supernatant was centrifuged for 10 min at 11400 g. Next, supernatant from this centrifugation was used for isolation of ER and pellet for isolation of mitochondria. The pellet was washed with 0.6 M mannitol; 10 mM Tris/Cl, pH 7.4, and the supernatant was centrifuged once at 17400 g and second time at 30000 g for 30 min. The final pellet contained ER membranes. Both, mitochondria and ER fractions, were resuspended in 10 mM Tris/HCl, pH 7.4. Purity of isolated fractions was checked with immunoblot of specific antibodies against relevant proteins localized to specific organelles.

### Phospholipid analysis

Lipids were extracted from isolated fractions by modified Folch extraction using chloroform/methanol/HCl (60:30:0.26; v/v) (Folch *et al*, 1957). The remaining contaminants were removed by additional washing steps of the organic phase with 0.1 % MgCl_2_ solution (w/v). Individual phospholipids were separated by thin layer chromatography on Silica gel 60 plates (Merck) using chloroform/methanol/acetic acid developing solvent (65:25:8; v/v/v). Phospholipids were visualized on thin layer chromatography plates by staining with iodine vapor, scraped off and quantified as described (Broekhuyse, 1968), with slight modifications. Briefly, thin layer chromatography plate was moistened with ultra-pure water and each phospholipid spot was scraped off, put into a glass tube and dried in an oven at 105 °C. After cooling, 200 μl of H_2_SO_4_/HClO_4_ (9:1; v/v) were added and the samples were incubated at 200 °C for 30 min. The samples were cooled down at room temperature and then incubated with 4.8 ml of 0.26 % (NH_4_)_6_Mo_7_O_24_.4 H_2_O (w/v) /ANSA (16 % K_2_S_2_O_5_; 0.252 % C_10_H_9_NO_4_S; 0.5 % Na_2_SO_3_; w/v) at a ratio of 250:11 (v/v) at 105 °C for other 30 min. After incubation, samples were cooled down, vortexed, and the silicagel was gently spun down at 500 g for 2 min. Phosphate (P_i_) of phospholipids reacts with Ammonium Heptamolybdate forming molybdenum blue, the concentration of which was assessed by absorption spectrometry at 830 nm. Pgc1 activity was measured as described previously (Simocková *et al*, 2008): *in vitro* reaction of final volume 240 μl contained 40 μl of 0.3 M Tris/HCl, pH 7.4; 20 μl of NBD-PG (0.8 μg) in 0.1% Triton X-100, and isolated subcellular fraction with defined amounts of proteins (LD fraction - 20 μg of proteins, other fractions - 175 μg of proteins) or a defined volume of liposomes (adequate for 200 μg of proteins of relevant cellular fraction, or 150 μl of buffered synthetic liposomes, see below). The mixture was incubated at 30 °C for 40 min. The reaction was stopped by the addition of chloroform/methanol/HCl (2 ml; 100:100:0.6 v/v/v) and 1 ml of water. After the separation, the organic phase was dried under a stream of nitrogen. NBD-labeled lipids were separated by one-dimensional thin layer chromatography for 17 min using chloroform/methanol/28% ammonium hydroxide developing solvent (65:35:5; v/v/v). Separated NBD-lipids were scanned with a TLC Scanner (Camag) in the fluorescence mode at wavelength 460 nm and phospholipase activity of Pgc1 was determined as a fraction of NBD-DAG, product of NBD-PG hydrolysis, in total NBD-lipids. Mixtures of LD (isolated from WT, *pgc1*Δ or *pgc1*Δ+*PGC1* cells) with various membrane bilayers (isolated from mitochondria or ER from *pgc1*Δ cells) were incubated at 30 °C on a shaker for 30 min.

### Preparation of liposomes

Liposomes were prepared either from phospholipids extracted from organelle fractions or from synthetic phospholipids (Avanti Polar Lipids). After evaporation of organic solvents under the stream of nitrogen, lipids extracted from isolated mitochondrial or ER fractions were buffered with 10 mM Tris/HCl, pH 7.4, and left at room temperature for one hour. Then, the suspension was sonicated (5 minutes) until it became transparent. In preparation of synthetic liposomes, 1,2-dipalmitoyl-*sn*-glycero-3-phosphocholine and 1-palmitoyl-2-oleoyl-*sn*-3-phosphoethanolamine (3:1; Avanti Polar Lipids) were dissolved in chloroform, which was evaporated under the stream of nitrogen. 10 mM Tris/HCl, pH 7.4 was added to a final concentration of phospholipids at 6.5 mM. Buffered lipids were incubated at 60 °C with occasional vortexing for 3-4 h. Then, the suspension was sonicated (5 minutes) until it became transparent. The final concentration of phospholipids was 4.1 mM.

### Immunoblotting

Proteins obtained from cell homogenates (100 μg) were separated on 12% SDS polyacrylamide gels and blotted onto a nitrocellulose membrane. The membrane was blocked using 5% dry milk in TBS buffer (50 mM Tris/HCl, pH 8.0; 150 mM NaCl; 0.05% Tween 20 (v/v)) overnight. Then, the membrane was immunostained using the following antibodies: polyclonal rabbit anti-GFP (Abcam, Cambridge, UK; dilution 1:1000); and rabbit anti-Por1 (1:1000). Visualization of secondary anti-rabbit (Sigma-Aldrich) antibodies was done using the Amersham ECL+ kit (GE Healthcare Life Sciences).

### Miscellaneous

Analysis of fatty acids, RNA isolation, RT-qPCR analysis (Pokorná *et al*, 2015) and isolation of LD fraction (Grillitsch *et al*, 2011) were performed as described previously. Statistical comparisons were carried out by one-way analysis of variance using SigmaPlot 12.5 software (Systat Software Inc.). All graphs show the mean ± SEM.

## Acknowledgements

We thank Marta Kostolanska, Petronela Melicherova and Katarina Nagyova for technical help. We are also grateful to Ivan Hapala for careful reading of the manuscript and stimulating comments. The work was supported by Scientific Grant Agency of the Ministry of Education, Science, Research and Sport of the Slovak Republic and the Slovak Academy of Sciences 2/0168/14 and 2/0165/18, the Slovak Research and Development Agency contracts APVV-15-0654, AS CR & SAV Joint Project SAV-18-25 and Slovak Academy of Sciences (SAS-MOSTJRP 2016/4). It is a result of the “Advanced Bioimaging of Living Tissues” project, reg. n. CZ.2.16/3.1.00/21527, which was financed from the budget of the European Regional Development Fund and public budgets of the Czech Republic through the Operational Programme Prague – Competitiveness. Günter Daum was supported by the Austrian Science Fund (FWF).

## Author contributions

MB and DK designed and performed most of the experiments and analyzed the data. JM and TA designed and performed the FRAP experiments. JM and PV performed fluorescence microscopy localizations. MB and JM wrote the manuscript with input from GD. All authors discussed the results.

---

**Figure EV1.**
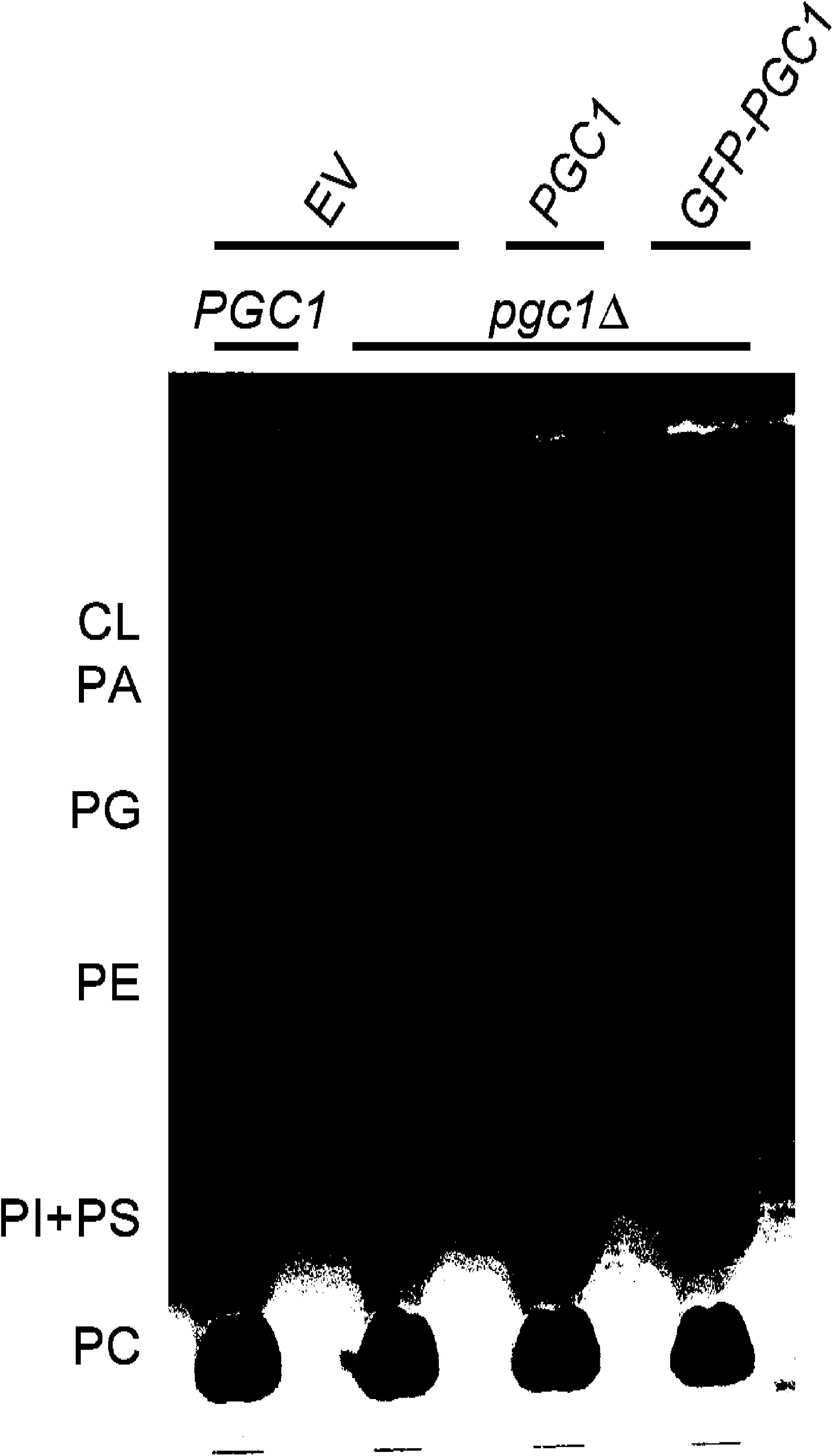
Expression of GFP-Pgc1 protein complements phenotype of *pgc1*Δ mutant. Phospholipids isolated from the yeast strains (WT+EV, pgc1Δ+EV, *pgc1*Δ+*PGC1, pgc1*Δ+*GFP-PGC1*) grown on SMD I- media without uracil were separated on thin layer chromatography and visualized as described in Materials and Methods. Note the absence of PG accumulation in the strains transformed with vector expressing *PGC1* or *GFP-PGC1* gene. PC, phosphatidylcholine; PI, phosphatidylinositol; PS, phosphatidylserine; PE, phosphatidylethanolamine; PG, phosphatidylglycerol; PA, phosphatidic acid; CL, cardiolipin; EV, empty vector.

**Figure EV2.**
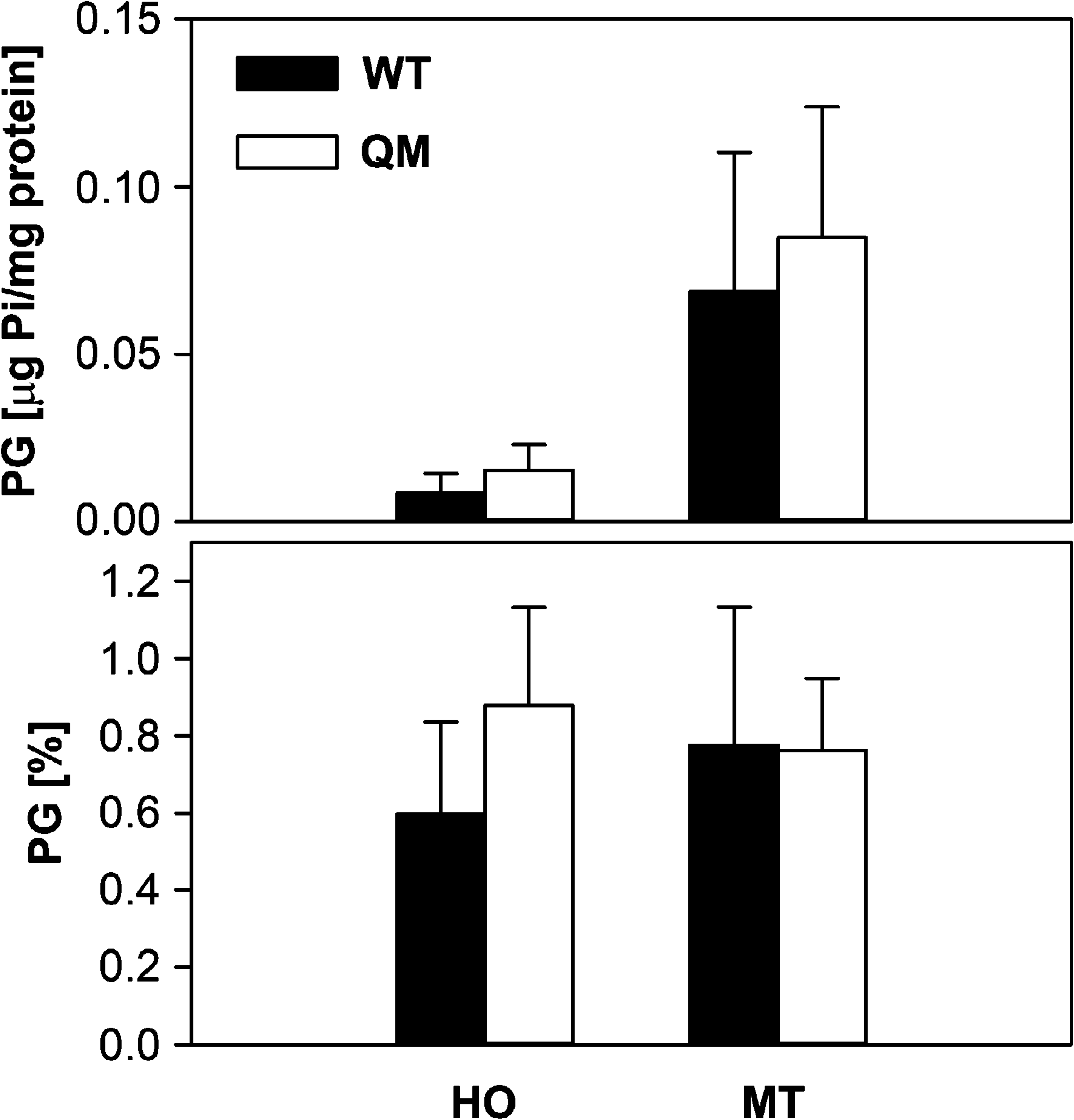
LD are dispensable for PG degradation. Wild type and QM cells were cultivated in SMD I- media for 24 h. Phospholipids were extracted from cell homogenate (HO) and isolated mitochondrial fraction (MT), separated by thin layer chromatography and visualised by iodine vapor. PG levels were quantified as described in Materials and Methods. Mean values of at least five independent experiments ± SEM are presented.

**Figure EV3.**
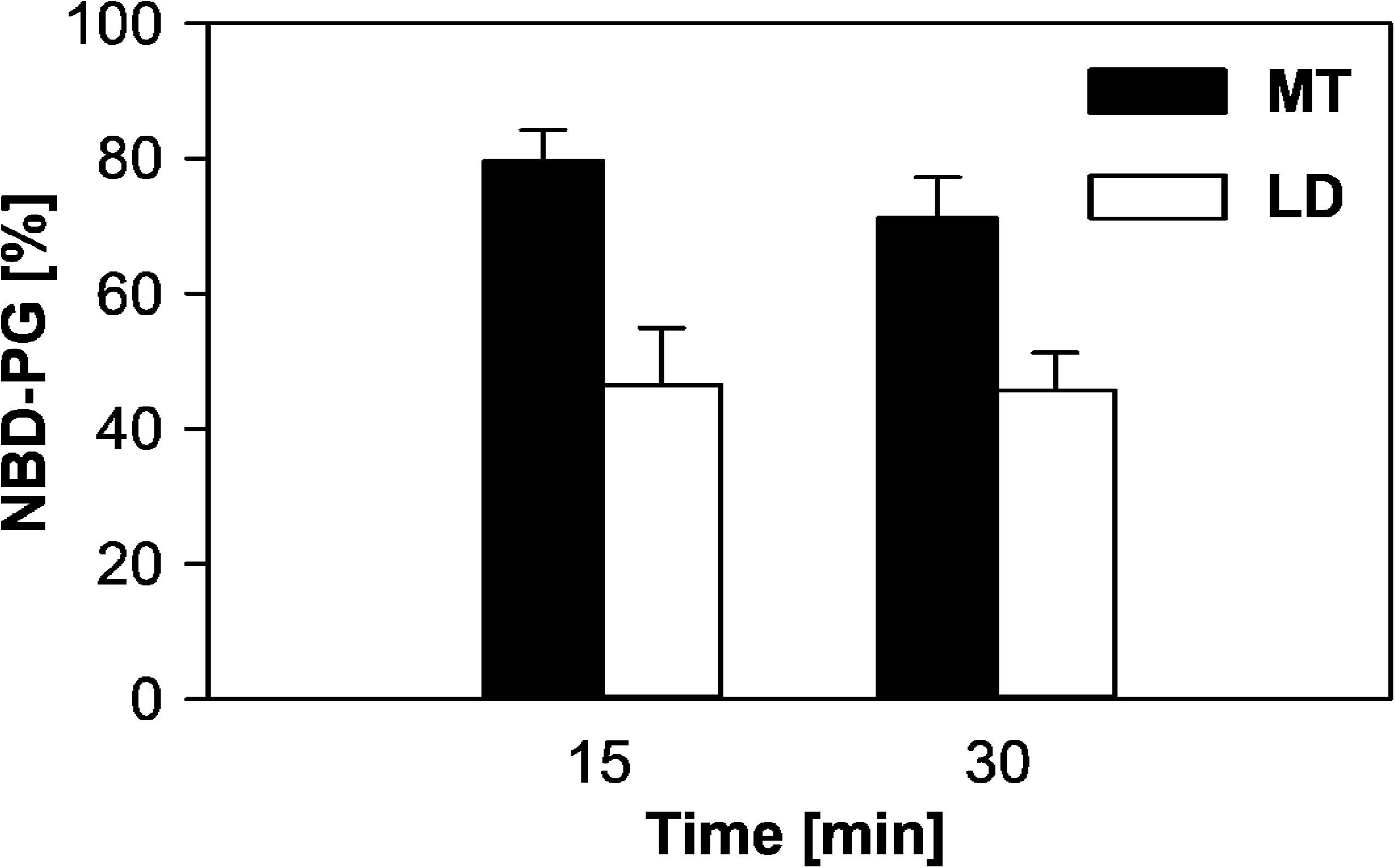
Fluorescently labelled PG is incorporated both to lipid mono- and bilayers. Incorporation of NBD-PG into the lipid monolayer at the surface of LD (empty bars) and mitochondrial membranes (full bars) was tested. In both cases, 0.8 μg NBD-PG was solved in 0.1 % Triton-X100 and added to suspension containing distinct amount of isolated wild type cellular fraction (20 μg of LD proteins, 175 μg of mitochondrial proteins) in 10 mM Tris/HCl, pH 7.4. Final volume of the mixture was 240 μΙ Mixtures were incubated at 30 °C on shaker for 15 or 30 minutes. Organelles were either floated (LD) or pelleted (mitochondria) by centrifugation and washed. Extracted phospholipids were laid on thin layer chromatography plate and fluorescent NBD-PG was visualized by TLC scanner (Camag). Mean values of two independent experiments ± SEM are presented.

**Figure EV4.**
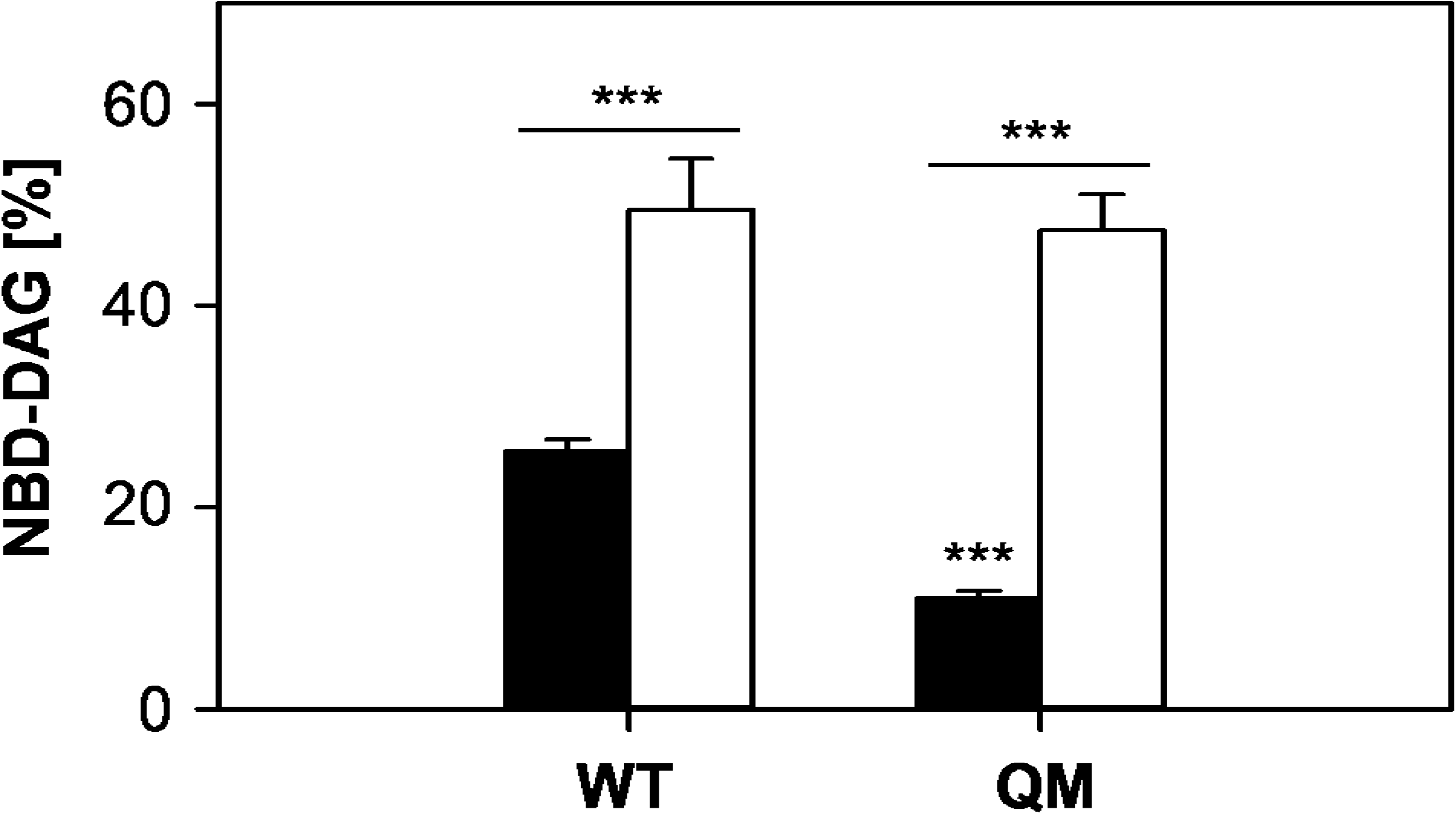
Presence of intact membranes enhances phospholipase activity of Pgc1. *In vitro* activities of Pgc1 in the wild type and QM strains were measured in cell homogenates prepared either by Dounce homogenizer as described in Materials and Methods (empty bars) or by glass beads (full bars). Glass bead homogenization was performed as follows: washed cells were suspended in 1 ml of 10 mM Tris/HCl, pH 7.4 and disrupted with glass beads (diameter 0.4 mm) in FastPrep (MP Biomedicals), for 2 × 45 s at the highest speed with 5 min cooling on ice between the runs. Mean values of at least four independent experiments ± SEM are presented. Statistically significant (see Materials and Methods for details) differences are marked by asterisks: *** - p < 0.001.

